# Simple and efficient heterologous expression of necrosis-inducing effectors using the model plant *Nicotiana benthamiana*

**DOI:** 10.1101/2021.05.02.442377

**Authors:** Bayantes Dagvadorj, Peter S. Solomon

## Abstract

Plant fungal pathogens cause devastating diseases on cereal plants and threaten global food security. During infection, these pathogens secrete proteinaceous effectors that promote disease. Some of these effectors from necrotrophic plant pathogens induce a cell death response (necrosis), which facilitates pathogen growth *in planta*. Characterisation of these effectors typically requires heterologous expression and microbial expression systems such as bacteria and yeast are the predominantly used. However, microbial expression systems often require optimization for any given effector and are, in general, not suitable for effectors involving cysteine bridges and posttranslational modifications for activity. Here, we describe a simple and efficient method for expressing such effectors in the model plant *Nicotiana benthamiana*. Briefly, an effector protein is transiently expressed and secreted into the apoplast of *N. benthamiana* by Agrobacterium-mediated infiltration. Two-to-three days subsequent to agroinfiltration, the apoplast from the infiltrated leaves is extracted and can be directly used for phenotyping on host plants. The efficacy of this approach was demonstrated by expressing the ToxA, Tox3 and Tox1 necrosis-inducing effectors from *Parastagonospora nodorum*. All three effectors produced in *N. benthamiana* were capable of inducing necrosis in wheat lines, and two of three showed visible bands on Coomassie-stained gel. These data suggest that *N. benthamiana-*agroinfiltration system is a feasible tool to obtain fungal effectors, especially those that require disulfide bonds and posttranslational modifications. Furthermore, due to the low number of proteins typically observed in the apoplast (compared to intracellular), this simple and high-throughput approach circumvents the requirement to lyse cells and further purify the target proteins that is required in other heterologous systems. Because of its simplicity and potential for high-throughput, this method is highly amenable to the phenotyping of candidate protein effectors on host plants.

## Introduction

Cereal production is susceptible to severe yield losses due to plant pathogens. During infection, plant pathogens secrete a repertoire of proteins called effectors into the apoplast or cytoplasm of the host cells to modulate plant processes and promote infection (Dodds and Rathjen 2010). Whilst these effectors underpin key disease mechanisms, many can be recognised by the plant, and the outcome of this interaction is broadly dependent on the the lifecycle of the specific pathogen. In biotrophic pathogens, effectors can be recognised by specific host receptors, triggering programmed cell death at the site of infection that results in plant resistance against these pathogens (Dodds and Rathjen 2010). In contrast, recent findings suggest that some necrotrophic pathogens, such as *Parastagonospora nodorum,* hijack these host resistance pathways for their own benefit (McDonald and Solomon 2018). Necrotrophic effectors (NEs) induce cell death only in the presence of host dominant susceptibility genes that strikingly resemble typical defence receptors (Lorang et al. 2012; Faris et al. 2010; Shi et al. 2016). Recognition, and the subsequent host cell death, is favoured by necrotrophic pathogens leading to plant susceptibility instead of resistance (Faris and Friesen 2020).

Despite recognising the important role effectors have in mediating disease, there are several challenges in understanding their function. These challenges are exemplified in studying effectors secreted by pathogens of non-model plants that lack genetic amenability. For example, most monocotyledon plants, such as economically important wheat and rice, are not compatible with the agroinfiltration method that allows transient expression of proteins in plants. Although the stable transformation has recently become possible in some of these plant species, including wheat, the approach is slow and cumbersome. To counter this, researchers have often turned to expressing their effectors of interest in model plant systems, such as *N. benthamiana*, as many effectors and their host cognate receptors are active in such systems (Vleeshouwers et al. 2008; Van Der Hoorn et al. 2000; Ma et al. 2012; Chen et al. 2017). In these cases, the transient expression of the effector with the corresponding host receptor is often sufficient to produce a cell-death phenotype that allows functional characterisation of their interaction (e.g. determining required amino acids for recognition etc). However, this approach does not appear to work for NEs (particularly from monocotyledon cereal pathogens) and their dominant susceptibility genes and co-expression in model systems such as *N. benthamiana* does not lead to a visible cell death. Therefore, this lack of a visible phenotype precludes this co-expression approach from being used to study these interactions (as it can be with many biotrophic effectors). Consequently, as an alternative approach, NEs are often expressed using heterologous systems for characterisation and subsequent phenotyping on host plants (Zhang et al. 2017; Outram et al. 2020; Sung et al. 2021; Liu et al. 2012, 2009). Almost exclusively, NEs have been expressed by using microbial expression systems such as *Escherichia coli* and *Pichia Pastoris*.

Many known and predicted effectors are small cysteine-rich (SCR) proteins (Saunders et al. 2012; Duplessis et al. 2011; Sperschneider et al. 2015; Stergiopoulos and De Wit 2009). It has been demonstrated that the cysteines in many of these effectors form disulfide bonds providing stability for these proteins in the apoplast (Saunders et al. 2012). For instance, the Tox3 and Tox1 effectors from *P. nodorum* contains six and sixteen cysteines respectively, and at least one disulfide bond is required for their necrosis-inducing activity (Liu et al. 2012, 2009; Outram et al. 2020). Similarly, *Cladosporium fulvum* effector Avr2 has four disulfide bonds that are essential for its function of inhibiting cysteine protease Rcr3 in tomato plants (Rooney et al. 2005). Moreover, some effectors such as ToxA and Tox3 appear to contain pro-domains subsequent to their secretion signals which are thought to be important for folding and are processed prior to secretion from the fungi (Outram et al. 2020; Tuori et al. 2000; Liu et al. 2009). Recently, Outram et al., (2020) reported that the pro-domain of Tox3 was cleaved by KEX2 protease that lead to a processed protein with demonstrably higher activity indicating that some necrotrophic effectors require posttranslational modification and processing to become fully active. However, because of these requirements, such proteins are difficult to express using a bacterial system due to the absence of eukaryotic cell organelles necessary for disulfide bond formation and posttranslational modifications. Lobstein *et al.* (2012) developed a new bacterial strain, *E. coli* SHuffle, which favors cytoplasmic disulfide bond formations. This new strain allowed researchers to express effectors harbouring disulfide bonds in soluble forms (Zhang et al. 2017; Outram et al. 2020). For example, one study demonstrated that the *E. coli* SHuffle system was successful in producing four out of five SCRs that were trialled (Zhang et al. 2017). Subsequent functional studies on three of the effectors and showed they were biologically active (Zhang et al. 2017). However, yields were often low limiting subsequent functional characterisation. Another approach to express SCR effectors has been using secretion pathways in eukaryotic cells such as yeast. Using this method, the SCR effectors SnTox1 and SnTox3 were successfully expressed at low levels in secreted forms and showed activity in wheat lines harboring corresponding susceptibility genes (Liu et al. 2009, 2012).

However, in microbial expression systems, various physical parameters need to be adjusted to achieve optimal expression conditions that can vary among different strains and target proteins. For example, in the *E. coli* SHuffle system, the effect of temperature and inducer concentration were specific to a given protein when tested on seven different proteins requiring disulfide bonds (Lobstein et al. 2012). Likewise, in the *P. pastoris* system, methanol is the primary carbon source, and its concentration must be closely monitored depending on the rate of utilization by the cells; less methanol decreases productivity, whereas too much is toxic to cells (Karbalaei et al. 2020). In addition to these examples, many other parameters in both *E. coli* and *P. pastoris* most likely need optimization for every new target protein (Lobstein et al. 2012; Karbalaei et al. 2020), which can be time-consuming and laborious. This can be a significant bottleneck for laboratories who do not have substantial protein expertise or are not experienced with microbial expression systems.

Here, we describe a simple and high-throughput method for heterologous expression of cysteine-rich effectors using the secretion pathway of model plant *N. benthamiana*. We demonstrate the efficacy of this method by expressing three unrelated necrosis-inducing effectors (that differ in their cysteine content) from *P. nodorum* and subsequent investigation of their activity on the host plant. Our data show that this plant expression approach can be used for rapid on-host functional screening of candidate necrotrophic effectors, especially those that require disulfide bonds and posttranslational modifications.

## Materials / Methods

### Plant material and growth conditions

*Triticum aestivum* cultivars Grandin, Corack, Calingari, and BG261 were grown in a controlled growth chamber with 250 μE light intensity, 85% relative humidity, and photoperiod of 16-h light at 20°C/8-h dark at 12°C. *Nicotiana benthamiana* plants were grown at 22°C with 16-h light/8-h dark cycle in a growth room.

### Cloning the effector proteins

The signal peptide sequences (SPs) of the effector proteins were predicted using SignalP-5.0 (Almagro Armenteros et al. 2019). The predicted SPs were replaced with *Nicotiana tabacum* PR-1 signal peptide (NtPR1sp) to promote efficient secretion of the effectors in *N. benthamiana*. These effector constructs were synthesized as gBlocks (IDT, Integrated DNA Technology), which were cloned into pENTR/D-TOPO gateway entry vector. The entry vectors were sequence confirmed and recombined into the pB7FWG2 gateway destination vector carrying C-terminus tag (GFP) fusion. To prevent the tag fusion to the effectors, we kept the original stop codons in the recombinant effector constructs, which allow the effectors to be expressed without the protein tag. The sequence confirmed destination vectors were transformed by electroporation into *Agrobacterium tumefaciens* (GV3101, pMP90) for agro-infiltration assays.

### Agro-infiltration assays

Agro-infiltration assays were carried out as described previously (Dagvadorj et al. 2017). Briefly, *A. tumefaciens* carrying the effector constructs were streaked on LB-agar plates with antibiotics (Spectinomycin 100 μg/ml, Gentamycin 20 μg/ml and Rifampicin 100 μg/ml) and grown for 2-3 days at 28°C. The bacterial cells on the plate were collected by scraping with 1 ml pipette tip, transferred to new 1.5 ml microcentrifuge tube and resuspended in 1 ml distilled water. The cells were centrifuged at 4000 rpm for 5 min at room temperature. The cell pellet was washed in 1 ml fresh Agro-induction buffer (AI; 10 mM MES pH 5.6, 10 mM MgCl_2_) and centrifuged again under the same conditions. Finally, the pellet was suspended in 1 ml AI buffer, and the cell suspension was diluted with AI buffer until the optical density (OD) of the cell suspension was adjusted to 0.2 for infiltration experiments. To enhance transient expression level, the cell suspensions were co-infiltrated with *A. tumefaciens* containing p19 RNA silencing suppressor construct (OD of 0.1). *N. benthamiana* 4- to 6- week of age were agro-infiltrated, and the leaves were harvested at 2 days post-agroinfiltration for apoplast wash fluid extraction.

### Apoplast washing fluid extraction

Apoplast washing fluid (AWF) was extracted as previously described with minor modifications (O’Leary et al. 2014). *N. benthamiana* leaves expressing the effector constructs were detached and submerged in a beaker with Milli-Q water. The beaker was placed into a desiccator and a vacuum of −80 kPa applied for 1 min. By gradually releasing the vacuum, the air within the apoplastic space was infiltrated with the water, and the infiltrated area appeared dark translucent by naked eye. This process was repeated until the whole leaf was infiltrated. Then, the leaves were wiped gently with clean tissue to remove the surface water and then, 3-4 leaves sandwiched together in parafilm sheets, rolled and placed into a 20 ml syringe, in a 50 mL Falcon tube. To isolate AWF, the Falcon tube was centrifuged at 500 g for 10 min at 4°C. The collected AWF (400-500 μL/per leaf) was transferred to microcentrifuge tubes and centrifuged again at 17,000 g for 5 min at 4°C to remove cell debris and other insoluble materials. The supernatant was transferred to a fresh tube and used immediately or stored at −80 °C.

### MS for ToxA and Tox3 protein identification

For peptide mapping of the ToxA and Tox3 proteins, AWF samples were separated on a 15% SDS-PAGE gel and stained with Coomassie blue. The proteins of interest were excised and subjected to in-gel trypsin digestion as previously described (Sung et al. 2021). Subsequent mass spectrometry data were analysed using Max-Quant software, v.1.6.0.8 (Tyanova et al. 2016). Tryptic peptides were mapped to the sequence of ToxA and Tox3 by using the trypsin enzyme digestion in semi-specific mode.

### AWF infiltration into wheat leaves

AWF samples were infiltrated using a needleless syringe into the second leaf of 2-week old wheat cultivars until the infiltration zone reached 3-5 cm. The wheat leaves were checked for the induction of necrosis from one day post-infiltration (dpi) and recorded at 3 dpi.

## Results

To trial the expression of SCR proteins in *N. benthamiana*, the ToxA, Tox3 and Tox1 effector proteins from *P. nodorum* were chosen (Table 1). ToxA, Tox3 and Tox1 induce necrosis on wheat lines harbouring the susceptibility genes *Tsn1*, *Snn3* and *Snn1*, respectively. The mature forms of ToxA, Tox3 and Tox1 each contain two, six and sixteen cysteine residues respectively, and their ability to induce necrosis is reliant on disulphide bond formation (Liu et al. 2012; Zhang et al. 2017; Sarma et al. 2005). In addition, ToxA and Tox3 require posttranslational modifications such as processing of pro-domain to become fully active (Outram et al. 2020; Tuori et al. 2000).

**Table 1.** Analysis of *P. nodorum* effectors used in this study.

| SCR effector | Accession number | Number of cysteines | Predicted signal peptide by SignalP 5.0 | Reference |
| --- | --- | --- | --- | --- |
| ToxA | AGE15694.1 | 2 | 1-16 | (Friesen et al. 2006) |
| Tox3 | ACR78113.1 | 6 | 1-20 | (Liu et al. 2009) |
| Tox1 | JN791682.1 | 16 | 1-17 | (Liu et al. 2012) |

To secrete the effector proteins to the apoplast of *N. benthamiana*, all three effectors were cloned with the *Nicotiana tabacum* PR-1 signal peptide (NtPR1sp; Ohshima et al. 1987) as shown in Figure 1a. This modification promotes the proteins to follow correct secretion pathway in *N. benthamiana* (Ma et al. 2015; Breen et al. 2016; Grosse-Holz et al. 2018). The effectors were expressed under control of CaMV 35S promoter. After two days post agro-infiltration, we isolated the apoplast washing fluid (AWF) from *N. benthamina* leaves expressing the effectors. The proteins within extracted AWFs were resolved by SDS-PAGE for analysis (Figure 1b). Although no observable band was detected for Tox1, the AWF containing ToxA and Tox3 showed bands within the size ranges of 12-14 kDa and 17-18 kDa, respectively. These sizes are similar to the respective sizes of mature ToxA and Tox3 identified from *P. nodorum* culture filtrates, respectively. To confirm the identity of the observed proteins, the bands were excised from the gel and subjected to tryptic digest MS. As expected, ‘Band1’ and ‘Band2’ in Figure 1b were positively identified as ToxA and Tox3, respectively (Figure 1c). Our analysis of the samples revealed that pro-domain of ToxA and Tox3 could not be detected, whereas C-terminal region of ToxA and Tox3 appeared to be intact (Figure 1c). Together, these results suggest that mature forms of ToxA and Tox3 were present in the apoplast of *N. benthamiana*.

**Figure 1.**
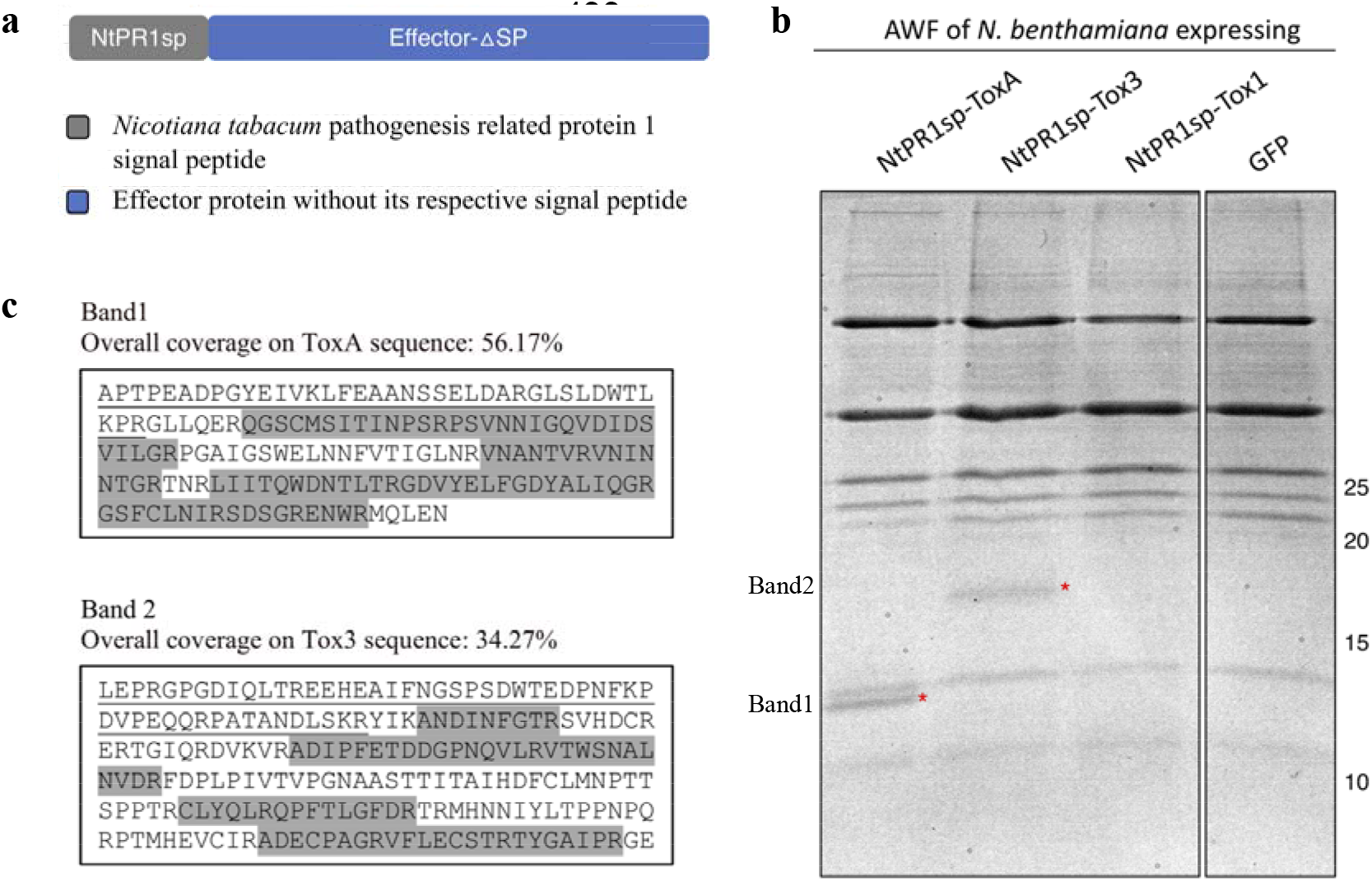
SCR effector proteins expression in *Nicotiana benthamiana*. (**a**) The schematic view of effector constructs used in this study. (**b**) Coomassie brillaint blue stained sodium dodecylsulfate-polyacrylamide gel electrophoresis (SDS-PAGE) analysis of apoplast washing fluid (AWF) isolated from *N. benthamiana* expressing various constructs. Astericks indicate the expected size of the matured ToxA and Tox3 proteins. AWF from *N. benthamiana* expressing GFP was used for background control. (**c**) Tryptic in-gel digest of ToxA and Tox3 from AWF of *N. benthamiana*. Tryptic peptides recovered after in-gel digest of corresponding ToxA and Tox3 bands, mapped to the sequence of ToxA and Tox3, respectively, are shown in grey. Predicted pro-domain sequences are underlined. Peptides were analysed by MaxQuant v1.6.8.0 specifying digestion mode as “Trypsin/P”.

To test biological activity of the effectors, the AWFs containing each of ToxA, Tox3, and Tox1 were infiltrated into wheat cultivars carrying *Tsn1*, *Snn3* and *Snn1* susceptibility genes (Grandin, Corack, and Calingari), respectively (Figure 2). Our results showed that the ToxA, Tox3 and Tox1 AWFs were able to induce necrosis in a genotype specific manner. Although we could not detect Tox1 band in protein gel analysis, we did observe chlorosis and necrosis development in the *Snn1* wheat cultivar (Calingari) infiltrated with AWF with Tox1 (Figure 2). A slight chlorotic response was also evident on the *snn1* cultivar Corack upon infiltration of the Tox1 AWF although a minor non-specific reaction for this effector has been previously noted when purified from *E. coli* (Zhang et al. 2017). Collectively, these results suggest that *N. benthamiana* expressed and secreted ToxA, Tox3 and Tox1 are biologically active.

**Figure 2.**
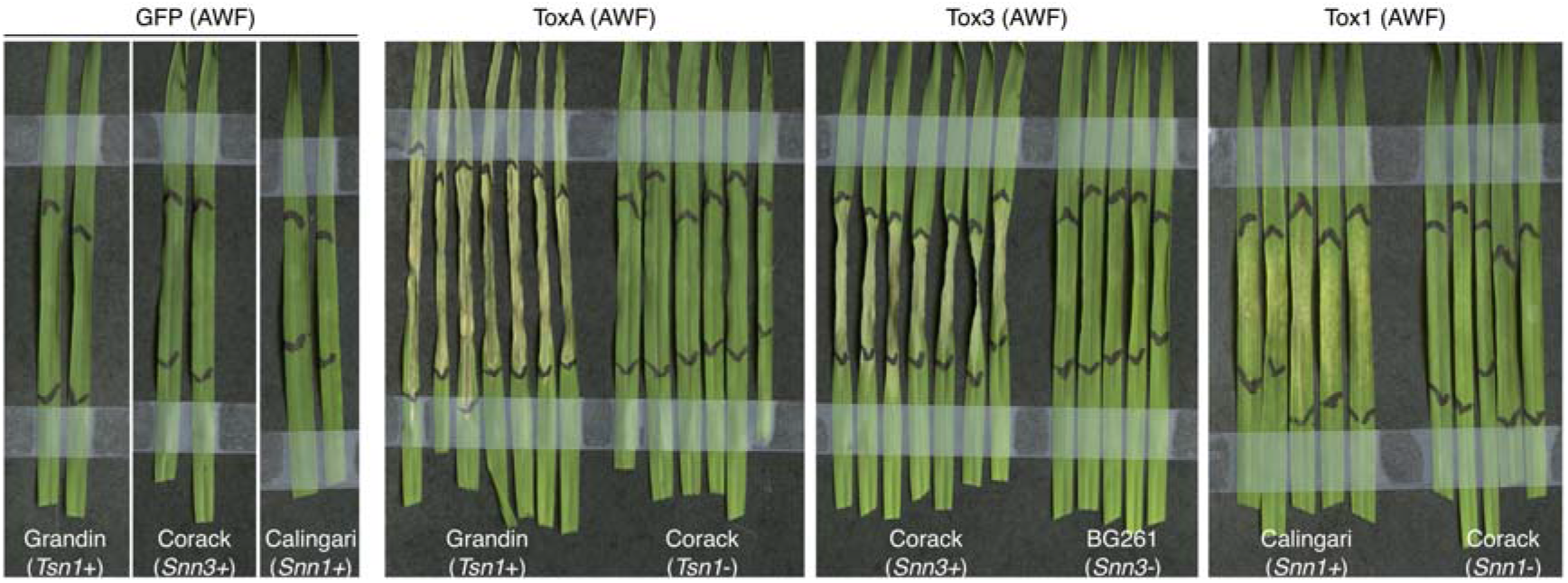
Apoplast washing fluid (AWF) containing the secreted ToxA, Tox3 and Tox1 trigger necrosis in wheat cultivars carrying the susceptibilty genes Grandin (*Tsn1*), Corack (*Snn3*) and Calingari (*Snn1*), respectively. Wheat cultivars Corack (*tsn1, snn1*) and BG261 (*snn3, snn1*) were used for genotype specificity controls, and AWF from *N. benthamiana* expressing GFP was used for AWF background control. The leaves were harvested and recorded three days after AWF infiltration.

## Discussion

In this study, we showed that fungal SCR effectors can be expressed and processed the using *N. benthamiana*-agroinfiltration system. Three unrelated SCR effectors, differing in their cysteine content, were successfully produced as active forms. In comparison to other methods currently used in the field, the approach we presented here is simple and efficient, which may facilitate the identification of putative NEs by screening for cell-death phenotype on host plants. This method may also be an alternative for researchers who struggle with expressing SCR effectors using microbial expression systems. In line with this, a recent study showed that using agroinfiltration, the apoplast targeted NLP2 (necrosis and ethylene-inducing peptide-like protein) was successfully expressed and induced necrosis in legume plants including field pea, faba bean and lentil; but not in chickpea (Debler et al. 2021). This further corroborates that plants can serve as a heterologous expression system to produce biologically active fungal effectors in secreted form.

This approach has several advantages compared with existing methods used to express necrotrophic effectors. Firstly, the plant signal peptide allows proteins to be trafficked through secretion pathways such as the endoplasmic reticulum and Golgi facilitating the formation of disulfide bonds and other posttranslational modifications. This facilitates the correct folding of SCR proteins are therefore more likely to be soluble compared to cytoplasmic-expressed proteins. Secondly, the apoplast can be separated mechanically from the plant cells, which enables the resolution of the mature proteins in the apoplast from the unprocessed forms inside the cell. This is not the case in cytosolically expressed proteins if crude extract used for the functional studies. Thirdly, the apoplast of the plant cell contains significantly less proteins and molecules than in the cytosol. Therefore, isolated apoplast wash fluid with target protein has much less background or contaminating proteins compared to proteins expressed in the cytosol for functional studies. Furthermore, the reduced protein content in the apoplast simplifies subsequent purification of the target protein if required. Although yeast eukaryotic systems can provide these advantages, various parameters influence the optimal expression, which can differ for any given protein. Thus, the optimization process is more challenging than compared with the *N. benthamiana*-agroinfiltration system.

However, the limitations of this method must be acknowledged. This approach may not be suitable for certain effectors, specifically those that trigger rapid cell-death in *N. benthamiana*. Several studies reported that effectors caused cell-death in *N. benthamiana* when expressed with their respective signal peptide (Kettles et al. 2017; Dagvadorj et al. 2017; Fang et al. 2016; Ma et al. 2015). For instance, eleven out of 119 putative effectors from *Ustilaginoidea virens* induced cell-death in *N. benthamiana*; however, the cause of the cell-death was unknown. In other studies, a group of effectors from *Zymoseptoria tritici* (Kettles et al. 2017), XEG1 effector from *Phytophthora sojae* (Ma et al. 2015), and PstSCR1 effector from *Puccinia striiformis* (Dagvadorj et al. 2017) triggered cell-death in *N. benthamiana,* which appear to be dependent on immune receptors such as NbSER3/NbBAK1. These results suggest that some effectors can activate basal immunity resulting the plant cell-death. For these types of effectors, non-plant systems may be more suitable.

Our study shows that *N. benthamiana*-agroinfiltration system can serve as an expression system for fungal cysteine-rich effectors. The expression of three unrelated effectors using this system displays production of high and biologically active effectors. This method can be upscaled and combined with a typical protein purification procedure to purify the protein of interest with little efforts. More importantly, this system applicable for expressing almost any protein (with some exceptions) resides in an extracellular space of the cell.

## Abbreviations

AWF: Apoplast washing fluid
NEs: Necrotrophic effectors
NtPR1sp: *Nicotiana tabacum* pathogenesis-related protein 1 signal peptide
SCR: small cysteine-rich

## Acknowledgements

This research was supported by the Australian Research Council (ARC) Discovery Project DP180102355. The authors thank Yi-Chang Sun for his technical assistance with the MS experiment.

## Notes

**Conflicts of interest** The authors declare that they have no conflicts of interest.

### Competing Interest Statement

The authors have declared no competing interest.

## References

Almagro Armenteros, J. J., Tsirigos, K. D., Sønderby, C. K., Petersen, T. N., Winther, O., Brunak, S., von Heijne, G., and Nielsen, H. 2019. SignalP 5.0 improves signal peptide predictions using deep neural networks. Nat. Biotechnol. 37:420–423

Breen, S., Williams, S. J., Winterberg, B., Kobe, B., and Solomon, P. S. 2016. Wheat PR-1 proteins are targeted by necrotrophic pathogen effector proteins. Plant J. 88:13–25

Chen, J., Upadhyaya, N. M., Ortiz, D., Sperschneider, J., Li, F., Bouton, C., Breen, S., Dong, C., Xu, B., Zhang, X., Mago, R., Newell, K., Xia, X., Bernoux, M., Taylor, J. M., Steffenson, B., Jin, Y., Zhang, P., Kanyuka, K., Figueroa, M., Ellis, J. G., Park, R. F., and Dodds, P. N. 2017. Loss of *AvrSr50* by somatic exchange in stem rust leads to virulence for *Sr50* resistance in wheat. Science. 358:1607–1610

Dagvadorj, B., Ozketen, A. C., Andac, A., Duggan, C., Bozkurt, T. O., and Akkaya, M. S. 2017. A *Puccinia striiformis* f. sp. *tritici* secreted protein activates plant immunity at the cell surface. Sci. Rep. 7:1141

Debler, J. W., Henares, B. M., and Lee, R. C. 2021. Agroinfiltration for transient gene expression and characterisation of fungal pathogen effectors in cool L season grain legume hosts. Plant Cell Rep.

Dodds, P. N., and Rathjen, J. P. 2010. Plant immunity: towards an integrated view of plant– pathogen interactions. Nat. Rev. Genet. 11:539–548

Duplessis, S., Cuomo, C. A., Lin, Y. C., Aerts, A., Tisserant, E., Veneault-Fourrey, C., Joly, D. L., Hacquard, S., Amselem, J., Cantarel, B. L., Chiu, R., Coutinho, P. M., Feau, N., Field, M., Frey, P., Gelhaye, E., Goldberg, J., Grabherr, M. G., Kodira, C. D., Kohler, A., Kues, U., Lindquist, E. A., Lucas, S. M., Mago, R., Mauceli, E., Morin, E., Murat, C., Pangilinan, J. L., Park, R., Pearson, M., Quesneville, H., Rouhier, N., Sakthikumar, S., Salamov, A. A., Schmutz, J., Selles, B., Shapiro, H., Tanguay, P., Tuskan, G. A., Henrissat, B., Van de Peer, Y., Rouze, P., Ellis, J. G., Dodds, P. N., Schein, J. E., Zhong, S., Hamelin, R. C., Grigoriev, I. V, Szabo, L. J., and Martin, F. 2011. From the Cover: Obligate biotrophy features unraveled by the genomic analysis of rust fungi. Proc Natl Acad Sci U S A. 108:9166–9171

Fang, A., Han, Y., Zhang, N., Zhang, M., Liu, L., Li, S., Lu, F., and Sun, W. 2016. Identification and Characterization of Plant Cell Death-Inducing Secreted Proteins From *Ustilaginoidea virens*. Mol. Plant. Microbe. Interact. 29:405–416

Faris, J. D., and Friesen, T. L. 2020. Plant genes hijacked by necrotrophic fungal pathogens. Curr. Opin. Plant Biol. 56:74–80

Faris, J. D., Zhang, Z., Lu, H., Lu, S., Reddy, L., Cloutier, S., Fellers, J. P., Meinhardt, S. W., Rasmussen, J. B., Xu, S. S., Oliver, R. P., Simons, K. J., and Friesen, T. L. 2010. A unique wheat disease resistance-like gene governs effector-triggered susceptibility to necrotrophic pathogens. Proc. Natl. Acad. Sci. 107:13544–13549

Friesen, T. L., Stukenbrock, E. H., Liu, Z., Meinhardt, S., Ling, H., Faris, J. D., Rasmussen, J. B., Solomon, P. S., McDonald, B. A., and Oliver, R. P. 2006. Emergence of a new disease as a result of interspecific virulence gene transfer. Nat. Genet. 38:953–956

Grosse-Holz, F., Madeira, L., Zahid, M. A., Songer, M., Kourelis, J., Fesenko, M., Ninck, S., Kaschani, F., Kaiser, M., and van der Hoorn, R. A. L. 2018. Three unrelated protease inhibitors enhance accumulation of pharmaceutical recombinant proteins in *Nicotiana benthamiana*. Plant Biotechnol. J. 16:1797–1810

Van Der Hoorn, R. A. L., Laurent, F., Roth, R., and De Wit, P. J. G. M. 2000. Agroinfiltration is a versatile tool that facilitates comparative analyses of *Avr9/Cf-9*-induced and *Avr4/Cf-4*-induced necrosis. Mol. Plant-Microbe Interact. 13:439–446

Karbalaei, M., Rezaee, S. A., and Farsiani, H. 2020. *Pichia pastoris*: A highly successful expression system for optimal synthesis of heterologous proteins. J. Cell. Physiol. 235:5867–5881

Kettles, G. J., Bayon, C., Canning, G., Rudd, J. J., and Kanyuka, K. 2017. Apoplastic recognition of multiple candidate effectors from the wheat pathogen *Zymoseptoria tritici* in the nonhost plant Nicotiana benthamiana. New Phytol. 213:338–350

Liu, Z., Faris, J. D., Oliver, R. P., Tan, K. C., Solomon, P. S., McDonald, M. C., McDonald, B. A., Nunez, A., Lu, S., Rasmussen, J. B., and Friesen, T. L. 2009. SnTox3 acts in effector triggered susceptibility to induce disease on wheat carrying the *Snn3* gene. PLoS Pathog. 5:e1000581

Liu, Z., Zhang, Z., Faris, J. D., Oliver, R. P., Syme, R., and Mcdonald, M. C. 2012. The cysteine rich necrotrophic effector SnTox1 produced by *Stagonospora nodorum* triggers susceptibility of wheat lines harboring Snn1. PLoS Pathog. 8:e1002467

Lobstein, J., Emrich, C. A., Jeans, C., Faulkner, M., Riggs, P., and Berkmen, M. 2012. Erratum to: SHuffle, a novel *Escherichia coli* protein expression strain capable of correctly folding disulfide bonded proteins in its cytoplasm. Microb. Cell Fact. 11:1–16

Lorang, J., Kidarsa, T., Bradford, C. S., Gilbert, B., Curtis, M., Tzeng, S. C., Maier, C. S., and Wolpert, T. J. 2012. Tricking the guard: Exploiting plant defense for disease susceptibility. Science. 338:659–662

Ma, L., Lukasik, E., Gawehns, F., and Takken, F. L. W. 2012. The use of Agroinfiltration for transient expression of plant resistance and fungal effector proteins in *Nicotiana benthamiana* leaves. Methods Mol. Biol. 835:61–74

Ma, Z., Song, T., Zhu, L., Ye, W., Wang, Y., Shao, Y., Dong, S., Zhang, Z., Dou, D., Zheng, X., Tyler, B. M., and Wang, Y. 2015. A *Phytophthora sojae* glycoside hydrolase 12 protein is a major virulence factor during soybean infection and is recognized as a PAMP. Plant Cell. 27:2057–2072

McDonald, M. C., and Solomon, P. S. 2018. Just the surface: advances in the discovery and characterization of necrotrophic wheat effectors. Curr. Opin. Microbiol. 46:14–18

O’Leary, B. M., Rico, A., McCraw, S., Fones, H. N., and Preston, G. M. 2014. The infiltration-centrifugation technique for extraction of apoplastic fluid from plant leaves using *Phaseolus vulgaris* as an example. J. Vis. Exp.:e52113

Ohshima, M., Matsuoka, M., Yamamoto, N., Tanaka, Y., Kano-Murakami, Y., Ozeki, Y., Kato, A., Harada, N., and Ohashi, Y. 1987. Nucleotide sequence of the PR-1 gene of *Nicotiana tabacum*. FEBS Lett. 225:243–246

Outram, M. A., Sung, Y.-C., Yu, D., Dagvadorj, B., Rima, S. A., Jones, D. A., Ericsson, D. J., Sperschneider, J., Solomon, P. S., Kobe, B., and Williams, S. J. 2020. The crystal structure of SnTox3 from the necrotrophic fungus *Parastagonospora nodorum* reveals a unique effector fold and insights into Kex2 protease processing of fungal effectors. bioRxiv.:2020.05.27.120113

Rooney, H. C., Van’t Klooster, J. W., van der Hoorn, R. A., Joosten, M. H., Jones, J. D., and de Wit, P. J. 2005. Cladosporium Avr2 inhibits tomato Rcr3 protease required for Cf-2-dependent disease resistance. Science. 308:1783–1786

Sarma, G. N., Manning, V. A., Ciuffetti, L. M., and Karplus, P. A. 2005. Structure of Ptr ToxA: An RGD-Containing Host-Selective Toxin from *Pyrenophora tritici-repentis*. 17:3190–3202

Saunders, D. G. O., Win, J., Cano, L. M., Szabo, L. J., Kamoun, S., and Raffaele, S. 2012. Using hierarchical clustering of secreted protein families to classify and rank candidate effectors of rust fungi. PLoS One. 7:e29847

Shi, G., Zhang, Z., Friesen, T. L., Raats, D., Fahima, T., Brueggeman, R. S., Lu, S., Trick, H. N., Liu, Z., Chao, W., Frenkel, Z., Xu, S. S., Rasmussen, J. B., and Faris, J. D. 2016. The hijacking of a receptor kinase-driven pathway by a wheat fungal pathogen leads to disease. Sci. Adv. 2:1–9

Sperschneider, J., Dodds, P. N., Gardiner, D. M., Manners, J. M., Singh, K. B., and Taylor, J. M. 2015. Advances and Challenges in Computational Prediction of Effectors from Plant Pathogenic Fungi. PLoS Pathog. 11:1–7

Stergiopoulos, I., and De Wit, P. J. G. M. 2009. Fungal effector proteins. Annu. Rev. Phytopathol. 47:233–263

Sung, Y., Outram, M. A., Breen, S., Wang, C., Dagvadorj, B., Winterberg, B., Kobe, B., Williams, S. J., and Solomon, P. S. 2021. PR1◻mediated defence via C◻terminal peptide release is targeted by a fungal pathogen effector. New Phytol. 229:3467–3480

Tuori, R. P., Wolpert, T. J., and Ciuffetti, L. M. 2000. Heterologous expression of functional Ptr ToxA. Mol. Plant-Microbe Interact. 13:456–464

Tyanova, S., Temu, T., and Cox, J. 2016. The MaxQuant computational platform for mass spectrometry-based shotgun proteomics. Nat. Protoc. 11:2301–2319

Vleeshouwers, V. G. A. A., Rietman, H., Krenek, P., Champouret, N., Oh, S., Wang, M., Bouwmeester, K., Vosman, B., Visser, R. G. F., Govers, F., Kamoun, S., and Vossen, E. A. G. Van Der. 2008. Effector Genomics Accelerates Discovery and Functional Profiling of Potato Disease Resistance and *Phytophthora Infestans* Avirulence Genes. 3:e2875

Zhang, X., Nguyen, N., Breen, S., Outram, M. A., Dodds, P. N., Kobe, B., Solomon, P. S., and Williams, S. J. 2017. Production of small cysteine-rich effector proteins in *Escherichia coli* for structural and functional studies. Mol. Plant Pathol. 18:141–151

